# LRRC8A anion channels modulate vasodilation via association with Myosin Phosphatase Rho Interacting Protein (MPRIP)

**DOI:** 10.1101/2023.03.08.531807

**Authors:** Hyehun Choi, Michael R. Miller, Hong-Ngan Nguyen, Jeffrey C. Rohrbough, Stephen R. Koch, Naoko Boatwright, Michael T. Yarboro, Rajan Sah, W. Hayes McDonald, J. Jeffrey Reese, Ryan J. Stark, Fred S. Lamb

**Author notes:** Corresponding author: Fred S. Lamb M.D. Ph.D., Vanderbilt University Medical Center, Department of Pediatrics, 2215 Garland Avenue, Light Hall-1055D, Nashville, TN 37232-3122.

## Abstract

**Background:** In vascular smooth muscle cells (VSMCs), LRRC8A volume regulated anion channels (VRACs) are activated by inflammatory and pro-contractile stimuli including tumor necrosis factor alpha (TNFα), angiotensin II and stretch. LRRC8A physically associates with NADPH oxidase 1 (Nox1) and supports its production of extracellular superoxide (O_2_^-•^).

**Methods and Results:** Mice lacking LRRC8A exclusively in VSMCs (Sm22α-Cre, KO) were used to assess the role of VRACs in TNFα signaling and vasomotor function. KO mesenteric vessels contracted normally to KCl and phenylephrine, but relaxation to acetylcholine (ACh) and sodium nitroprusside (SNP) was enhanced compared to wild type (WT). 48 hours of *ex vivo* exposure to TNFα (10ng/ml) markedly impaired dilation to ACh and SNP in WT but not KO vessels. VRAC blockade (carbenoxolone, CBX, 100 μM, 20 min) enhanced dilation of control rings and restored impaired dilation following TNFα exposure. Myogenic tone was absent in KO rings. LRRC8A immunoprecipitation followed by mass spectroscopy identified 35 proteins that interacted with LRRC8A. Pathway analysis revealed actin cytoskeletal regulation as the most closely associated function of these proteins. Among these proteins, the Myosin Phosphatase Rho-Interacting protein (MPRIP) links RhoA, MYPT1 and actin. LRRC8A-MPRIP co-localization was confirmed by confocal imaging of tagged proteins, Proximity Ligation Assays, and IP/western blots which revealed LRRC8A binding at the second Pleckstrin Homology domain of MPRIP. siLRRC8A or CBX treatment decreased RhoA activity in cultured VSMCs, and MYPT1 phosphorylation at T853 was reduced in KO mesenteries suggesting that reduced ROCK activity contributes to enhanced relaxation. MPRIP was a target of redox modification, becoming oxidized (sulfenylated) after TNFα exposure.

**Conclusions:** Interaction of Nox1/LRRC8A with MPRIP/RhoA/MYPT1/actin may allow redox regulation of the cytoskeleton and link Nox1 activation to both inflammation and vascular contractility.

## Introduction

Hypertension is a chronic vascular inflammatory disease. Serum levels of tumor necrosis factor alpha (TNFα)^1^, interleukin 1-beta (IL-1β)^2^ and the pro-inflammatory vasoconstrictor angiotensin II (AngII)^3^ are chronically elevated and inflammation correlates with severity of disease^4^. In vascular smooth muscle cells (VSMCs) these signaling pathways all depend upon activation of the Nox1 NADPH oxidase^5–7^. Furthermore, as blood pressure rises, increased wall tension promotes inflammation^8^ in a reactive oxygen species (ROS)-dependent manner.^9^ In resistance vessels, stretch activates the autoregulatory “myogenic” contractile response of VSMCs that helps to normalize local blood flow. This response is also Nox1^10^ and ROS-dependent^11^. While acutely adaptive, myogenic tone can drive further increases in vascular resistance and support a positive feedback loop of inflammation and hypertension that drives disease progression^12^. Thus, ROS that originate as Nox1-derived superoxide (O_2_^-•^) are critical mediators of both inflammatory and bio-mechanical signaling in VSMCs.

O_2_^-•^ can increase vascular tone by inactivation of nitric oxide (NO^•^). It has also been linked to activation of the RhoA small GTPase, the primary regulator of Rho-associated kinase (ROCK) which is hyperactivated in multiple forms of experimental hypertension^13^. ROCK regulates the calcium sensitivity of VSMC contraction by phosphorylating and inhibiting Myosin Phosphatase Target Subunit 1 (MYPT1), the regulatory component of the myosin light chain phosphatase (MLCP) complex. This increases MLC phosphorylation and promotes crossbridge cycling with actin.

Volume-regulated anion channels (VRACs) are composed of LRRC8 family proteins and are essential elements of a multi-protein signaling complex that also includes the type 1 TNFα receptor (TNFR1)^14^, Nox1 and the redox-sensitive MAPKKK5, also known as Apoptosis Signaling-related Kinase 1 (ASK1)^15^. Loss of either LRRC8A, the LRRC8 isoform that is required for all VRACs^16^ or its LRRC8C partner^17^ impaired Nox1 activation by TNFα. This disrupted multiple endpoints of downstream signaling in VSMCs including TNFα receptor endocytosis and Mitogen-Activated Protein Kinase (MAPK) and NF-κB activation. The basis for functional dependence of Nox1 on anion current is unknown. Extracellular scavenging of O_2_^-•^ by superoxide dismutase (SOD) interfered with TNFα signaling, while catalase was without effect^16^. Thus, O_2_^-•^ from Nox1 clearly plays a role beyond serving as a substrate for conversion to H_2_O_2_.

Since loss or block of LRRC8A channels so effectively protected cultured VSMCs from TNFα-induced inflammation^16^ we wanted to study this effect in intact blood vessels and created VSMC-specific LRRC8A knockout mice. KO mesenteric blood vessels had normal contractile responses but displayed remarkably enhanced vasodilation via multiple mechanisms, and completely lacked myogenic tone. TNFα signaling was completely abrogated. Immunoprecipitation of LRRC8A and mass spectroscopy identified Myosin Phosphatase Rho-Interacting Protein (MPRIP, a.k.a. M-RIP or p116Rip) as a binding partner for LRRC8A. MPRIP is a scaffolding protein that also binds to RhoA, MYPT1 and actin and is essential for regulation of the MLCP complex. Reduction of RhoA activity and ROCK-dependent MYPT1 phosphorylation in VSMCs with reduced LRRC8A protein levels or VRAC inhibition suggests that LRRC8A colocalizes TNFR1/Nox1-dependent O_2_^-•^ production with the regulatory machinery controlling MLCP activity. This supports the novel concept that inflammatory signaling impacts VSMC contractility via redox signaling by O_2_^-•^ that is dependent upon LRRC8A anion channels.

## Methods

Detailed methods are provided in the Supplemental Material. All animal studies complied with the *Guiding Principles in the Care and Use of Animals,* approved by the Vanderbilt University Institutional Animal Care and Use Committee.

## Results

### Vasomotor function in LRRC8A null vessels

Freshly isolated mesenteric vessel segments from WT and VSMC-specific LRRC8A KO mice were studied by wire myography. Compared to WT, KO vessels displayed indistinguishable contractile responses to depolarization by KCl or adrenergic receptor activation by phenylephrine (PE, Suppl Fig. 1A and B). Sustained contractile responses to KCl and PE are almost completely dependent on influx of extracellular calcium^18^. Following contraction with PE (10^-6^ M) KO vessels were more responsive to relaxation induced by acetylcholine (ACh, LogEC_50_, WT −6.2 ± 0.11 vs. KO −6.7 ± 0.07, *p* < 0.05) which elicits an endothelium-dependent relaxation response that is multi-factorial (Fig. 1A). An important component of the response is production of nitric oxide (NO^•^) by endothelial cells. To selectively evaluate responsiveness to NO^•^, relaxation in response to sodium nitroprusside (SNP) was evaluated (Fig, 1B). Relaxation to SNP was also enhanced in KO vessels (LogEC_50_, WT −8.1 ± 0.11 vs. KO −8.6 ± 0.13, *p* < 0.05).

**Figure 1.**
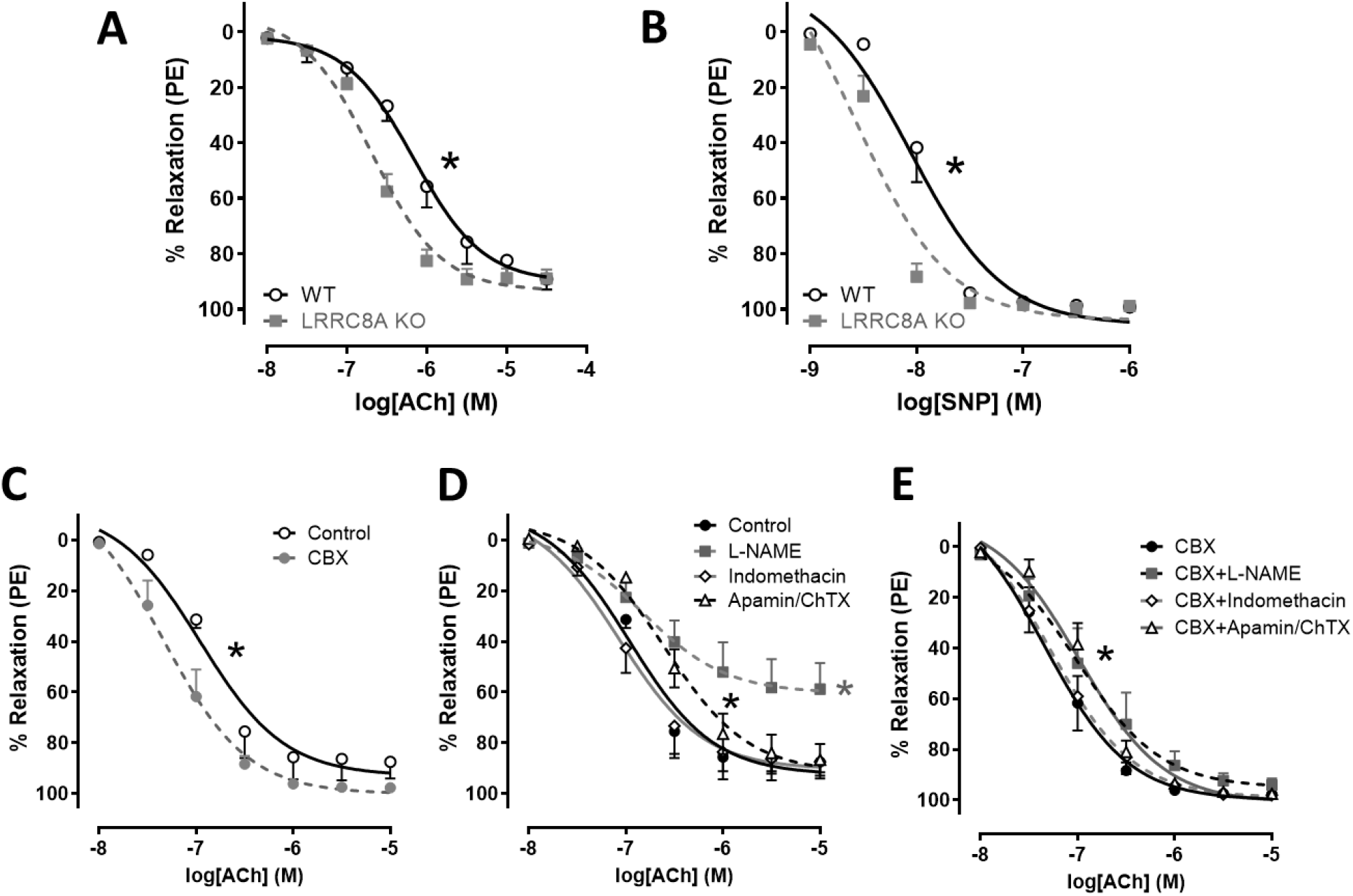
LRRC8A KO and carbenoxolone (CBX) augment vasodilation in mesenteric arteries. In freshly isolated mesenteric arteries, relaxation in response to **A)** acetylcholine (ACh, WT n = 6, KO n = 8) and **B)** sodium nitroprusside (SNP, WT n = 6, KO n = 5) was significantly augmented in KO compared to WT vessels. * Indicates *p* < 0.05 for EC_50_. **C)** 15 min incubation in CBX (100 μM) significantly enhanced relaxation responses to ACh. * Indicates *p* < 0.05 for EC_50_. **D)** Effect of inhibiting; NO synthase with L-NAME (100 μM, 30 min), prostaglandin synthesis with indomethacin (5 μM, 20 min), or potassium channels with Apamin (50 nM, 20 min) + Charybdotoxin (ChTX; 50 nM, 20 min), on ACh-induced relaxation under control conditions in WT vessels. **E)** Effect of the same drugs on the ACh response in WT vessels treated with CBX for 15 minutes. CBX-treatment enhances relaxation compared to untreated controls under all conditions tested. Black* indicates *p* < 0.05 Apamin/ChTX vs. Control (D) or CBX (E) for EC_50_. Gray* indicates *p* < 0.05 L-NAME vs. Control for Emax (n = 4 to 5).

**Supplemental Figure 1.**
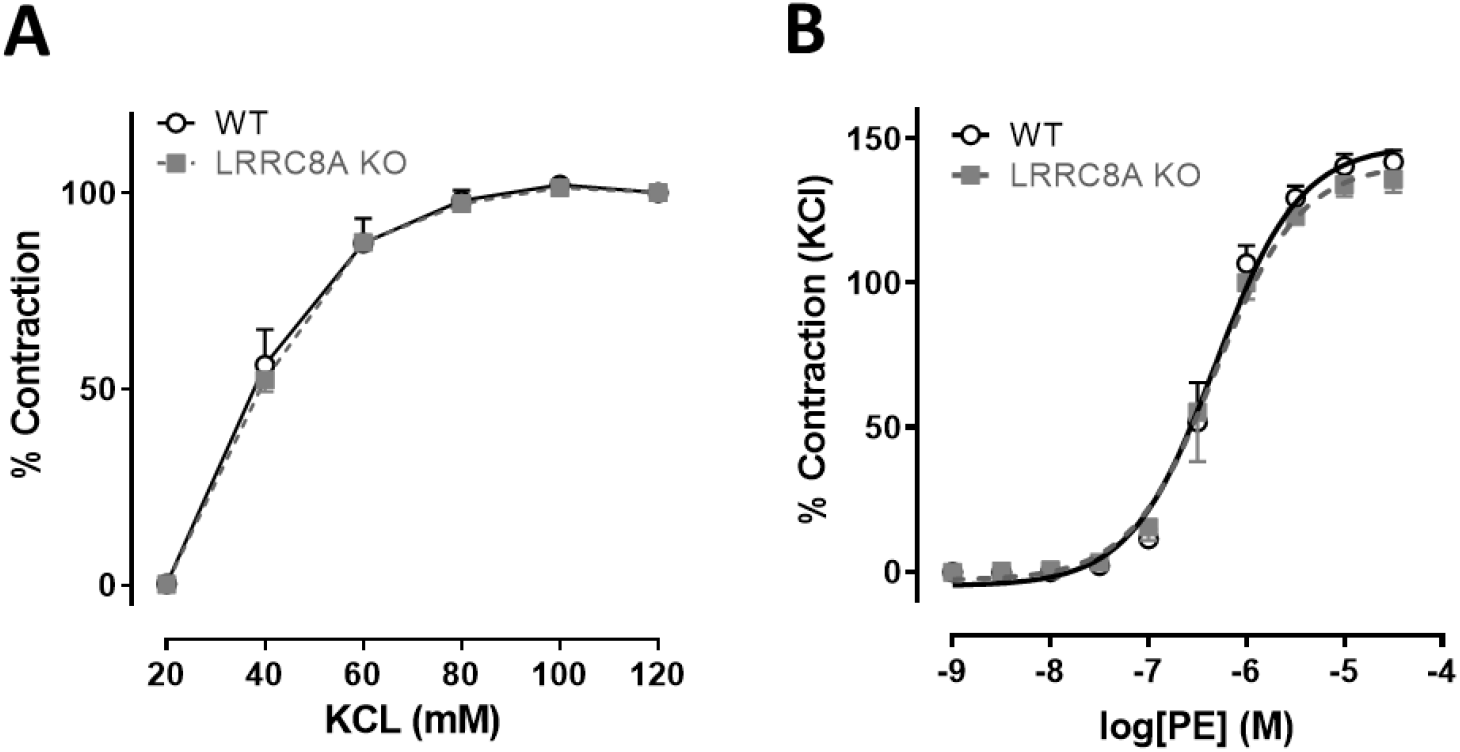
Contractile response of freshly isolated WT and LRRC8A KO mesenteric arteries. Responses to **A)** KCl (n = 3) and **B)** phenylephrine (PE, n = 4) were unaltered.

To determine if potentiation of vasodilation in LRRC8A KO vessels was related to loss of ion channel function, these experiments were repeated in WT vessel segments exposed to the VRAC inhibitor, carbenoxolone (CBX, 10^-4^ M) or vehicle (water) for 15 minutes prior to assessment of vasodilator function. As with KO vessels, CBX enhanced dilation to ACh (Fig. 1C and Suppl. Table 1). Because the response to ACh in mesenteric vessels reflects a summation of signaling via multiple dilator mechanisms, pharmacological tools were used to isolate those components (Fig. 1D and E and Suppl. Table 1). In control vessels (Fig. 1D) blocking prostaglandin production with indomethacin (5 μM) did not alter relaxation to ACh. Endothelium-dependent VSMC hyperpolarization can be inhibited by blockade of potassium channels with a mixture of apamin (50 nM) and charybdotoxin (ChTX, 50 nM). This combination decreased ACh sensitivity in control rings, making them approximately as sensitive as CBX-treated rings. NO^•^ production was inhibited using L-NAME (100 μM) and this profoundly impaired relaxation in control vessels where the maximal relaxation was significantly reduced. The response of CBX-treated rings to ACh (Fig. 1E) was also insensitive to indomethacin while apamin + ChTX caused a small rightward shift in the D/R curve but preserved a higher sensitivity compared to control. The most remarkable effect of CBX was on relaxation responses in L-NAME-treated rings. NO synthase inhibition still caused a small decrease in ACh sensitivity, but the ability of ACh to cause a full relaxation response was preserved. In summary, CBX-treated vessels relaxed significantly better than control under all conditions tested.

To assess the impact of VSMC LRRC8A KO on TNFα signaling, mesenteric vessel segments were incubated in tissue culture for 48hrs in the presence of TNFα (10ng/ml) or vehicle. As observed in freshly isolated tissue, vehicle-exposed cultured vessels from KO mice were more sensitive to both ACh (Fig. 2A) and SNP (Fig. 2B). TNFα markedly reduced relaxation in WT tissues in response to both ACh and SNP but had no effect on the responses of KO vessels. Impaired relaxation of WT rings following TNFα exposure could result from impaired endothelial dilator production and/or reduced VSMC responsiveness. Vessels from both groups would be expected to have intact endothelial cell responsiveness to TNFα and therefore a similar degree of endothelial dysfunction. The degree to which the endothelium contributes to this *in vitro* model of TNFα-induced inflammation is not discernable by these experiments. However, the clear lack of detrimental impact of TNFα on the response of KO vessels to SNP likely reflects a combination of impaired TNFα signaling and enhanced baseline vasodilator function in KO VSMCs.

**Figure 2.**
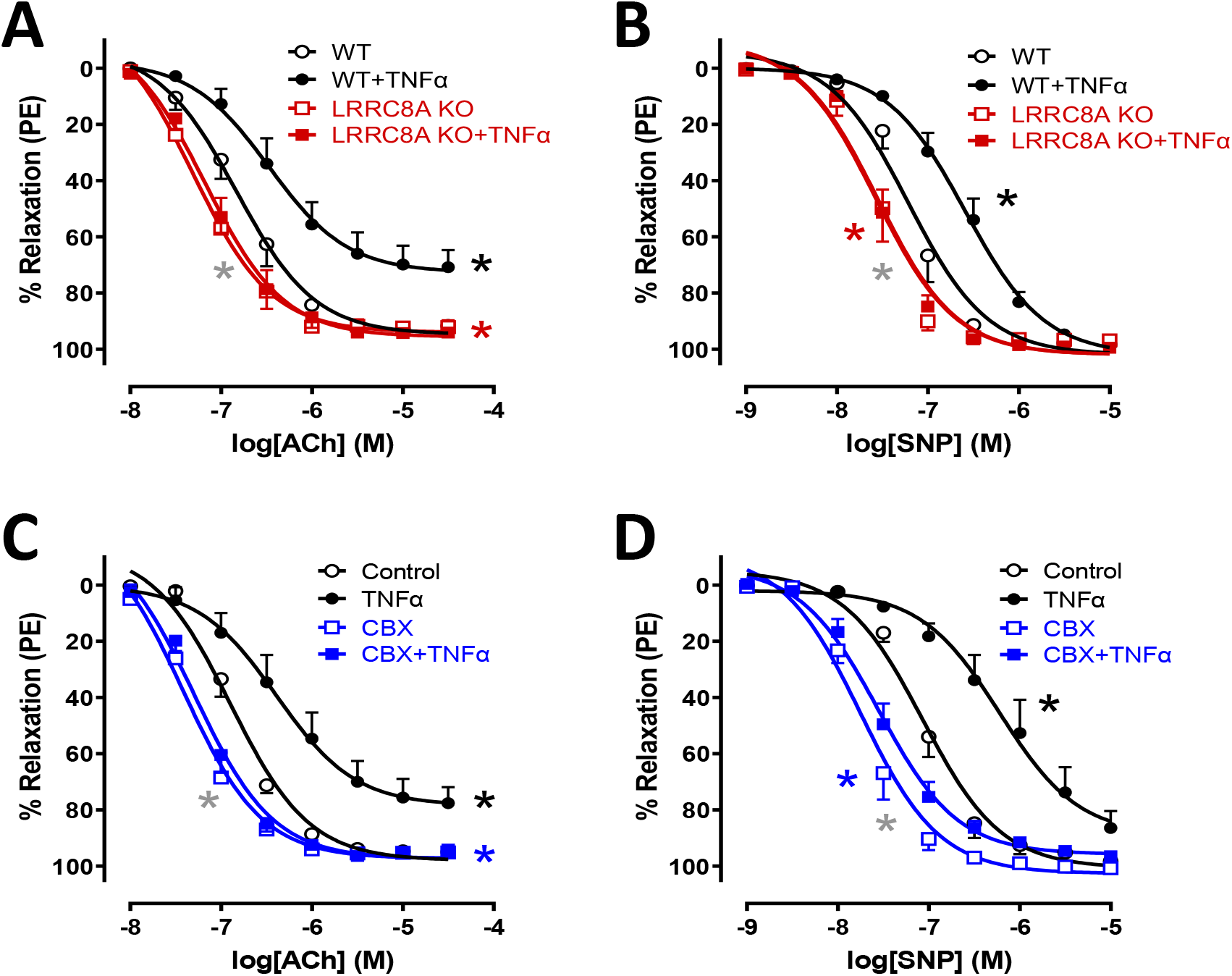
Disruption of LRRC8A function protects against TNFα-induced vascular dysfunction. WT and LRRC8A KO mesenteric vessels were incubated in tissue culture for 48 hrs in media ± TNFα (10ng/ml). Relaxation responses to **A)** ACh and **B)** SNP were assessed after constriction with PE. KO vessels relaxed significantly better than WT and were completely protected from impairment following TNFα exposure as seen in WT vessels (n = 6 - 8). **C and D)** Vessel segments from WT mice were exposed to TNFα for 48hrs and then following mounting for tension recording to CBX (100μM, 20min) or vehicle. CBX effectively reversed impaired dilation after TNFα exposure (n = 5). **A)** * Indicates *p* < 0.05 for WT vs. WT + TNFα (black*) for EC_50_ and E_max_, WT + TNFα vs. KO + TNFα (red*) for EC_50_ and E_max_, or WT vs. KO (gray*) for EC_50_. **B)** * Indicates *p* < 0.05 for WT vs. WT + TNFα (black*) for EC_50_, WT + TNFα vs. KO + TNFα (red*) for EC_50_, or WT vs. KO (gray*) for EC_50_. **C)** * Indicates *p* < 0.05 for Control vs. TNFα (black*) for EC_50_ and E_max_, TNFα vs. CBX+TNFα (blue*) for EC_50_ and E_max_, or Control vs. CBX (gray*) for EC_50_. **D)** * Indicates *p* < 0.05 for Control vs. TNFα (black*) for EC_50_, TNFα vs. CBX+TNFα (blue*) for EC_50_, or Control vs. CBX (gray*) for EC_50_.

TNFα-induced VSMC dysfunction after 48hrs could be related to VSMC “damage”, such as protein or lipid oxidation, or to changes in gene expression. Alternatively, TNFα could drive ongoing inactivation of NO^•^ by O_2_^-•^ or persistently stimulate O_2_^-•^-dependent signaling. These latter events would be expected to be rapidly reversible. We addressed this question by applying CBX treatment for only 20 min in the muscle bath following 48 hours of TNFα exposure. This brief period of VRAC inhibition by CBX provided protection from the adverse effects of cytokine exposure that was very similar to that provided by LRRC8A KO (Fig. 2C and D). Rescue of impaired vasodilation in response to both ACh and SNP by CBX suggests that VRAC inhibition disrupts an ongoing signal that interferes with VSMC relaxation.

Dilation of LRRC8A KO vessels might be enhanced by protection from NO^•^ inactivation. This would be consistent with our previous demonstration of reduced O_2_^-•^ production^16^. We reasoned that if this was the case, relaxation in response to vasodilator mechanisms that are downstream of, or unrelated to NO^•^, would be unaltered. We therefore assessed vasodilator responsiveness to agents that alter the function of three different kinases that phosphorylate MYPT1 and regulate MLCP activity. Following contraction with PE, WT and KO vessels were exposed to; 1) BAY60-2770, which directly activates soluble guanylyl cyclase, increases intracellular cyclic GMP, and activates protein kinase G (PKG), 2) Forskolin, which activates adenylyl cyclase, increases cyclic AMP levels and activates protein kinase A (PKA), or 3) Y-27632, which inhibits Rho-associated coiled-coil kinase (ROCK, Suppl Fig. 2). All three agents were fully effective vasodilators (~100% relaxation), and responsiveness was unaffected by TNFα exposure in either WT or KO tissues. This suggests that the mechanism(s) by which TNFα impairs vasodilation are upstream of MLCP and the contractile proteins. In contrast, the ED50 for all three drugs was significantly lower in KO compared to WT vessels. Thus, the

**Supplemental Figure 2.**
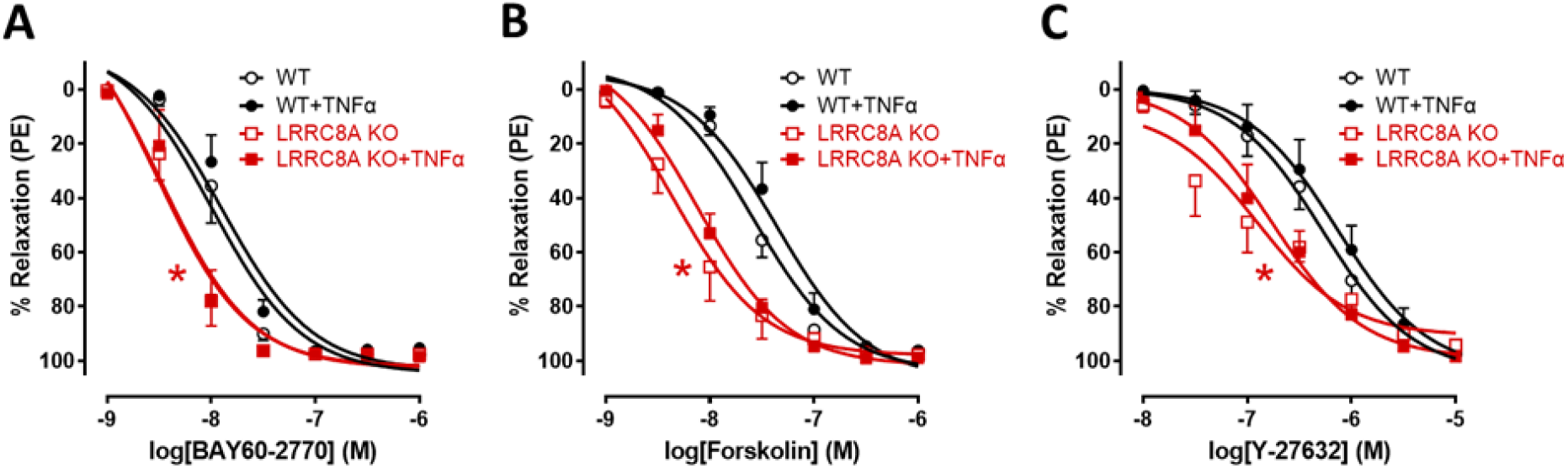
Loss of LRRC8A enhances relaxation by multiple mechanisms. Relaxation responses to activation of **A)** soluble guanylyl cyclase (BAY60-2770, n = 6 - 7), **B)** adenylate cyclase (Forskolin, n = 5 - 6) or **C)** inhibition of Rho kinase (Y-27632, n = 5 - 6). Knockout of LRRC8A in VSMCs significantly enhanced relaxation to all three agents. None of these responses were affected by TNFα exposure in WT or KO vessels. Red* indicates *p* < 0.05 for WT vs. KO. mechanism of enhanced relaxation in LRRC8A KO vessels is downstream of PKG, PKA and ROCK. This strongly implicated a fundamental change in MYPT1 regulation of MLCP.

**Suppl. Table 1.** LogEC_50_ and E_max_ values for ACh-induced responses in mesenteric arteries

| Agonist | LogEC<sub>50</sub> |  | E<sub>max</sub> |  |
| --- | --- | --- | --- | --- |
|  | Control | CBX | Control | CBX |
| Vehicle | -7.0±0.15 | -7.3±0.11* | 93±4.7 | 100±2.5 |
| L-NAME | -6.8±0.27 | -7.0±0.15 | 60±5.7‡ | 95±3.9 |
| Indomethacin | -7.1±0.15 | -7.3±0.12 | 90±4.4 | 99±3.0 |
| Apamin/ChTX | -6.6±0.11* | -7.0±0.08† | 92±4.3 | 101±2.3 |

Values are mean ± SEM for n experiments in each group. Relaxation induced by ACh and SNP was calculated relative to the maximal changes from the contraction produced by PE and is represented as percentage of relaxation. \**p*<0.05 vs Control in LogEC<sub>50</sub>; †*p*<0.05 vs CBX in LogEC<sub>50</sub>; ‡*p*<0.05 vs Control in E<sub>max</sub> (n = 4 to 5).

Myogenic tone is an innate response of VSMCs to physical stretch. Both Nox1 and ROS have been implicated in its regulation^10,19^. In addition, this response is associated with activation of RhoA/ROCK which enhances calcium sensitivity of the contractile process^20^. We reasoned that if loss of LRRC8A impacts vasodilation at the level of MLCP then myogenic tone might also be affected. Pressure myography was used to quantify the response of mesenteric vessel segments to increasing intraluminal pressure (20 - 140 mmHg, Fig. 3). A myogenic response was triggered in WT vessels beginning at ~80mmHg as demonstrated by differences in the diameter of vessels tested in normal vs. calcium-free buffer containing a potent vasodilator (papaverine, 10^-4^ M). The response of WT segments was comparable in magnitude to that previously reported in murine mesenteric vessels^21^ but no myogenic tone developed in KO vessels, which displayed exclusively passive dilation in response to increasing luminal pressure.

**Figure 3.**
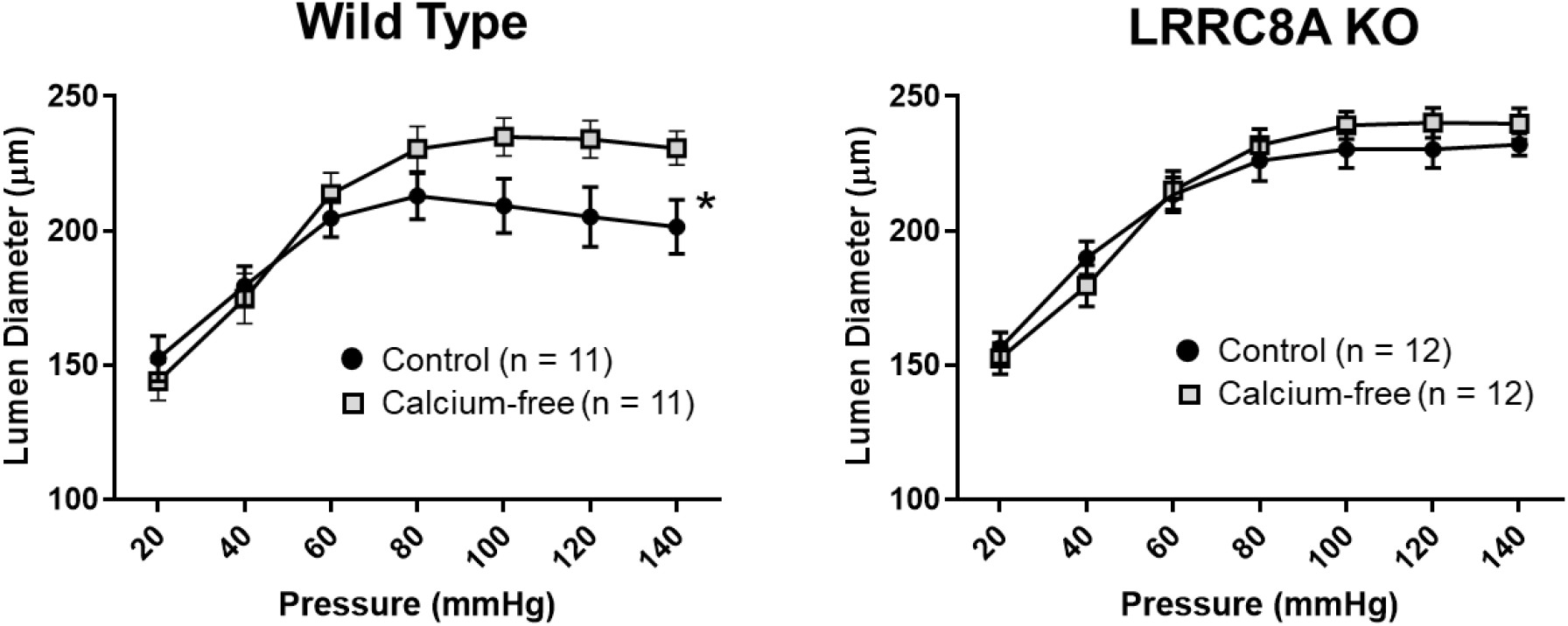
Myogenic tone is absent in LRRC8A KO mesenteric arteries. Pressure-induced constriction (the difference in diameter in normal vs. Ca^++^-free buffer + papaverine) was absent in mesenteric segments from LRRC8A VSMC KO mice. \**p* < 0.05 by 2-way ANOVA across all pressures (n = 11 and 12).

### Identification of LRRC8A-associated proteins

LRRC8 proteins have cytoplasmic C-terminal LRR domains that are involved in protein-protein interaction^22^ and seem likely to confer specific function to VRACs. To explore the mechanism by which LRRC8A impacts vasomotor function we immunoprecipitated proteins from WT and KO VSMCs with an anti-LRRC8A antibody and used mass spectroscopy to identify potential binding partners. Proteins pulled down in KO cells were used to control for non-specific binding. Over two combined experiments (Suppl. Table 2), 33 proteins (Fig 4A, columns 1 and 2) met our initial criteria for LRRC8A-association; at least 5 total reads, > 4 reads in the WT precipitate while either never being identified in the KO, or > 4X as many reads in WT cells. The most frequently identified proteins were LRRC8A and LRRC8C, and LRRC8D was also only found in WT cells, confirming that the pulldown was selective. Analysis of the interrelationship between the proteins was performed using STRING (Fig. 4C). The most associated biological process was “actin filament-based process” (red), the top molecular function was “actin binding” (blue), and the top cellular component was the “actin cytoskeleton” (green, Fig 4B). To further validate the comparison of WT and KO samples we analyzed the frequency of reads for proteins that are highly unlikely to be associated with a plasma membrane protein. This provided a control group of “background” reads that included annexins, collagens, histones, keratins, elongation factors, mitochondrial ATPase subunits, and heat shock, ribonuclear, RNA-binding and ribosomal proteins. Collectively, these reads were present in very similar abundance (WT 584, KO 597, Suppl. Table 2). This suggests a similar degree of non-specific pulldown from the two types of cells. We therefore added to our analysis proteins that had reads in high abundance, such as actin and myosin isoforms, but were identified more than twice as frequently in the WT. These proteins are listed in Fig. 4A, column 3. Including them significantly decreased the estimated false discovery rate for all 3 pathway categories (Fig. 4B, shaded data). The resulting proteinprotein interaction network is shown in Fig. 4C and is color coded for pathways of interest.

**Figure 4.**
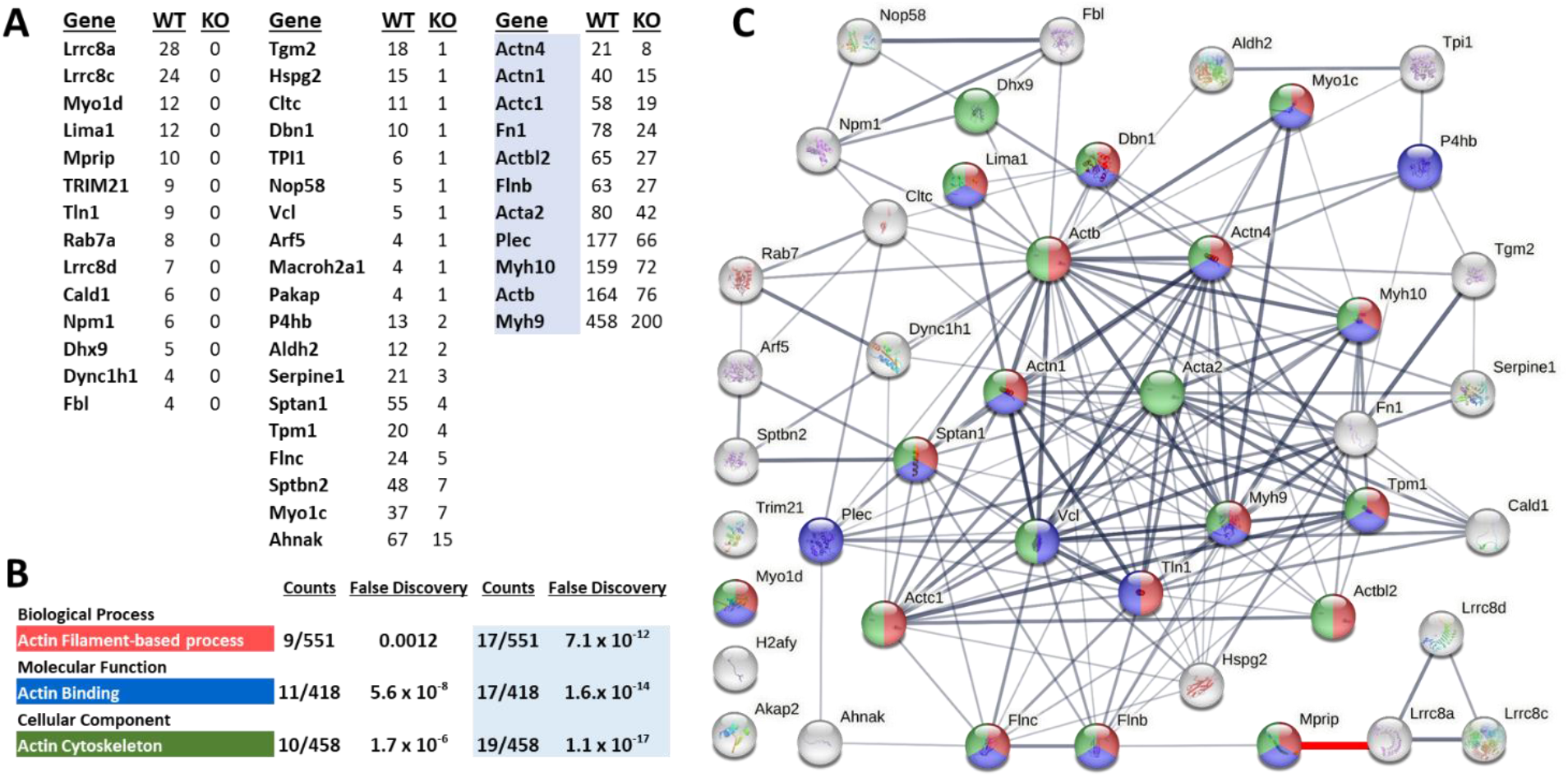
Identification of LRRC8A-associated proteins. **A)** Columns 1 and 2 list proteins that met initial criteria for selective immunoprecipitation by anti-LRRC8A as identified by mass spectroscopy (see Suppl. Table 2 for details). Column 3 proteins (blue highlight) met secondary criteria. **B)** Top result of pathway analysis for three categories of function, associated number of proteins detected for each pathway, and the false discovery rate for each observation. The last 2 columns provide data from a secondary analysis that incorporated the highlighted proteins in column 3. **C)** Association network generated by string-db.org. Proteins are color coded for their inclusion in the pathways listed in **(B)**. The thickness of connecting lines reflects the confidence of the association between proteins. A connection between MPRIP and LRRC8A has been added (red, lower right) to reflect the association demonstrated by the current work.

### MPRIP links LRRC8A with the cytoskeleton

Among the proteins that were uniquely pulled down in WT cells was MPRIP, a scaffolding protein known to colocalize the RhoA small GTPase with the MYPT1 regulatory subunit of MLCP and actin^23–25^. Knockdown of MPRIP disrupts RhoA regulation of MLCP^26^. We therefore sought to confirm association of LRRC8A with MPRIP and identify the site of LRRC8A binding. Western blotting of the WT and KO LRRC8A immunoprecipitates confirmed the presence of MPRIP in the WT but not the KO pulldown (Fig. 5A). Proximity Ligation Assays (PLA) were employed to further establish colocalization of the two proteins. A clear PLA signal was present in cultured WT but not KO VSMCs (Fig. 5B). Finally, full length MPRIP tagged with C-terminal meGFP was co-expressed with LRRC8A-mCherry in HEK293T cells (Fig. 5C). As previously demonstrated, LRRC8A localizes to the plasma membrane and to prominent intracellular vesicles^27^. MPRIP co-localized with LRRC8A at the membrane, particularly in areas where membrane extensions were forming, but was completely absent from intracellular vesicles. Collectively, these data confirm physical interaction between LRRC8A and MPRIP.

**Figure 5.**
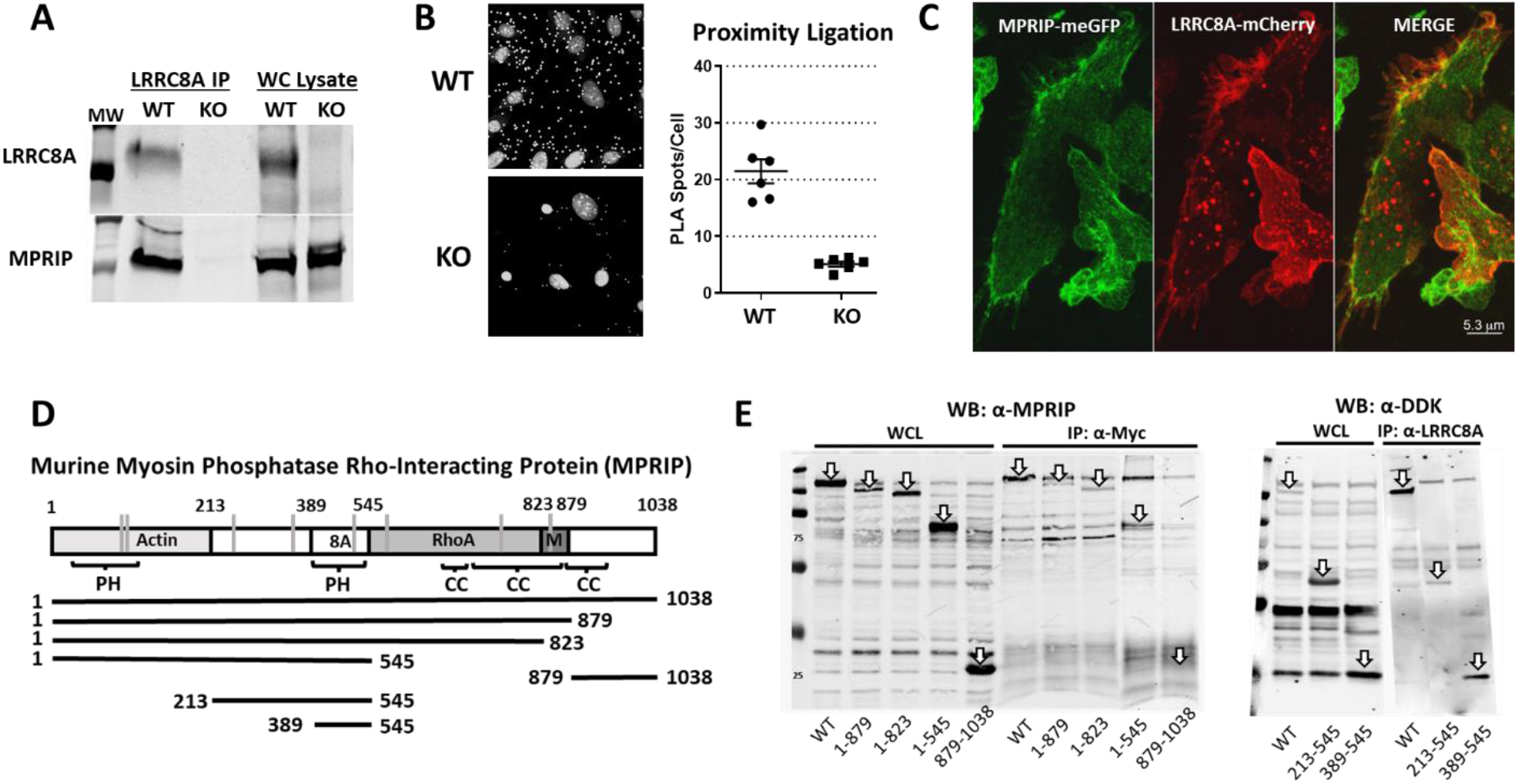
Association of LRRC8A and MPRIP. **A)** Immunoprecipitation (IP) of endogenous LRRC8A from VSMCs confirms the presence of LRRC8A in the whole cell (WC) lysate and precipitated protein from WT but not LRRC8A KO cells. MPRIP is present in the WC lysate from both genotypes but is only pulled down in LRRC8A WT cells. **B)** Proximity Ligation Assay (PLA) signal from α-MPRIP/α-LRRC8A in WT VSMCs. Small white spots represent PLA signal, the larger spots are nuclei. Counts of spots/nucleus from multiple fields of confluent WT and LRRC8A KO VSMCs (20x). PLA spots were smaller and significantly less numerous in KO cells. **C)** Confocal images of HEK293T cells co-expressing MPRIP-meGFP and LRRC8A-mCherry. Note colocalization at the plasma membrane, particularly at ruffled borders but not in LRRC8A-containing intracellular vesicles. **D)** Sites of protein binding to MPRIP. The first Plextrin Homology (PH) domain (aa 44-152) binds actin. C-terminal coiled-coil domains mediate interaction with RhoA and MYPT1 (M). Below the map are the constructs utilized to determine the site of MPRIP binding to LRRC8A (8A) was localized to aa 389-545. Vertical grey lines represent specific cysteines that can be oxidized based upon the Oxymouse database (Cys103, 120, 235, 361, 506, 571, 723, 830). Susceptible cysteines are present in the binding regions for all 4 protein partners. **E)** Immunoprecipitation of MPRIP-DDK fragments by full length LRRC8A-Myc in HEK293T cells expressing these constructs. Arrows identify overexpressed proteins, or the site where they would be expected to run (see IP: α-Myc) in whole cell lysates (WCL, left) and in immunoprecipitated protein (IP, right). All C-terminal MPRIP deletion mutants associate with LRRC8A, but the C-terminal MPRIP fragment 879-1038 does not associate with LRRC8A. The 213-545 peptide associated with LRRC8A as did the region between 389 and 545 which contains PH2 (see IP: α-LRRC8A).

The regions of MPRIP that bind its other partners have been previously identified. Actin associates with a Pleckstrin Homology (PH) domain in the N-terminal region (aa 1-213)^25^, RhoA binds to a region between aa 545 and 823^24^ and MYPT1 binding occurs adjacent to that (aa 824-879)^28^ as outlined in Fig. 5D. We created DDK-tagged C-terminal mutants of MPRIP and co-expressed these with full length myc-tagged LRRC8A in HEK293T cells. Immunoprecipitation with anti-Myc and western blotting with α-MPRIP revealed that LRRC8A associates with the region of MPRIP between the end of the actin binding region and the start of the RhoA binding sequence (aa 213-545). Full length MPRIP also associated with a peptide containing only aa 213-545, as well as a smaller subclone expressing only the region containing the second PH domain (aa 389-545) (Fig. 5E). These experiments define the region of MPRIP interaction with LRRC8A between aa389 and 545 that corresponds to a second PH domain.

### LRRC8A modulates RhoA/ROCK/MYPT1 function

The RhoA small GTPase is regulated by Nox1^29^ and by O_2_^-•30^. RhoA activates ROCK which phosphorylates MYPT1, inhibits phosphatase activity, and increases the calcium sensitivity the myosin light chain. MYPT1 phosphorylation by ROCK also impairs the ability of protein kinase A (PKA) and protein kinase G (PKG) to phosphorylate MYPT1 at adjacent sites which activates MLCP and triggers vasodilation. We compared RhoA activity in cultured WT VSMCs treated with scrambled or LRRC8A-targeted siRNA or exposed to CBX (100 μM) for 20 min (Fig. 6A). siLRRC8A significantly reduced LRRC8A protein abundance to 71 ± 3.2% of control (n = 6). To mimic conditions in our ring experiments, cells were exposed to PE (2 × 10^-6^ M) for 3 min followed by SNP (3 × 10^-8^ M) for an additional 3 min before lysis and assay of RhoA activity. LRRC8A knockdown was associated with significantly lower RhoA activity (siControl: 100 ± 2.0%, siLRRC8A: 89 ± 2.1% of control) as was CBX (Control: 100 ± 3.0%, CBX: 88 ± 4.2% of control). To determine if ROCK activity was affected, we assayed MYPT1 phosphorylation at T853 in acutely isolated, intact mesenteric beds. Mesenteries were exposed to the same conditions utilized for RhoA activity assays (PE followed by SNP), flash frozen and protein isolated for western blotting. Total MYPT1 abundance was similar between WT and KO, but phosphorylation at T853 was significantly reduced in KO mesenteries (Fig. 6B), consistent with diminished activity of RhoA/ROCK. Thus, consistent with the phenotypes of enhanced relaxation and impaired myogenic tone, loss of LRRC8A protein or function impaired Rho/ROCK activity.

**Figure 6.**
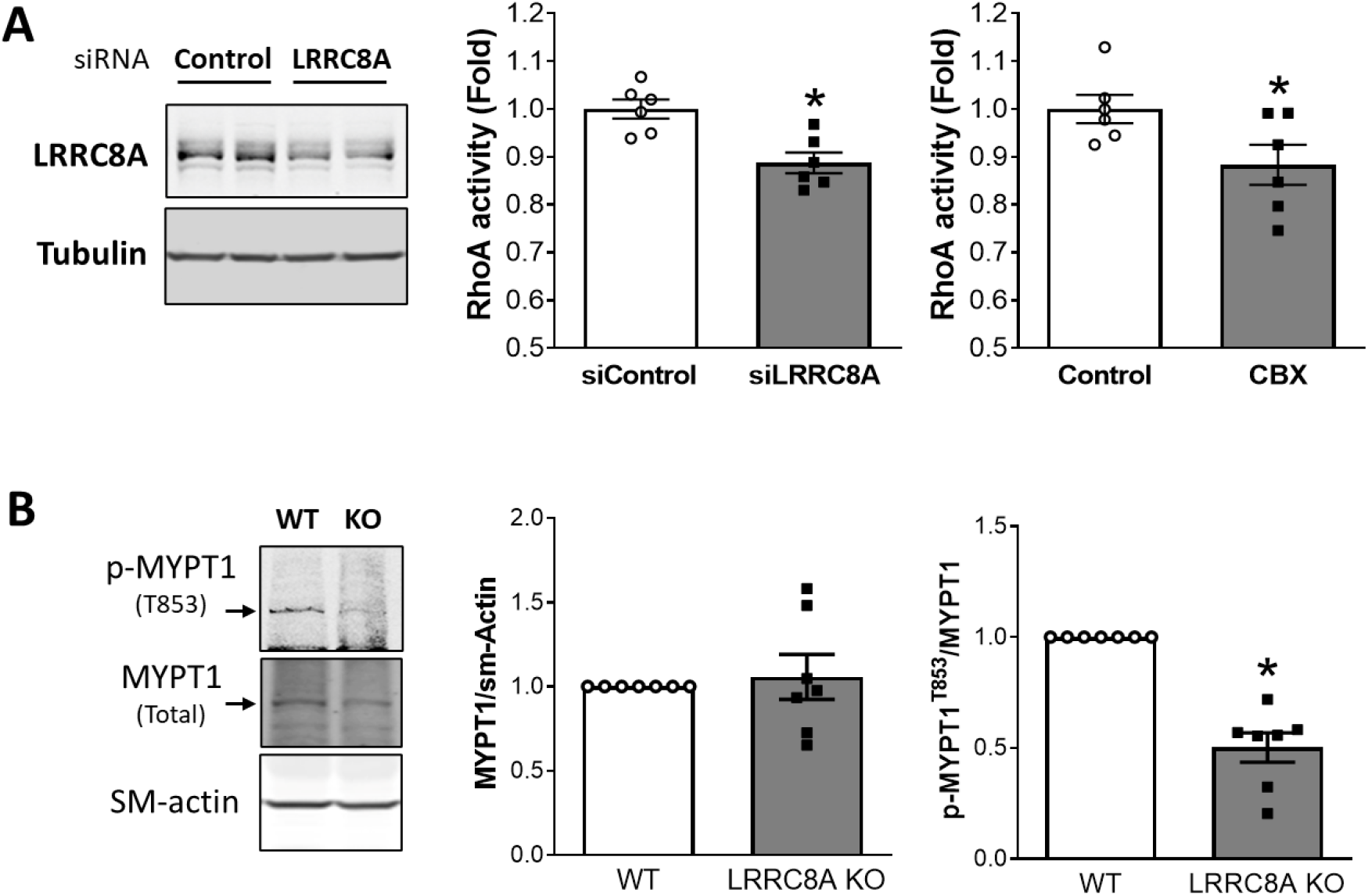
RhoA-signaling is LRRC8A-dependent. **A)** LRRC8A protein was significantly reduced in cultured mesenteric VSMCs by treatment with siLRRC8A. LRRRC8A knockdown and CBX treatment were associated with similar significant decreases in RhoA activity (n = 6). **B)** LRRC8A protein abundance is reduced but still detectable in intact mesenteries (fat, vessels and connective tissue) from KO mice (n = 7). Total MYPT1 protein is unaltered but phosphorylation at T853 (p-MYPT1^T853^) is significantly reduced in mesenteries from VSMC-specific LRRC8A KO mice. \**p* < 0.05 compared to siControl or WT.

To further investigate a role for MPRIP in LRRC8A-dependent signaling we utilized siMPRIP to reduce the abundance of the protein in cultured mesenteric and aortic VSMCs (Fig. 7A). MPRIP knockdown was significant in both cell types and was associated with a significant increase in the abundance of LRRC8A protein. This may represent an attempt by the cells to compensate for loss of a downstream signaling partner. In contrast, MPRIP expression was not altered in LRRC8A null VSMCs (Fig. 7B).

**Figure 7.**
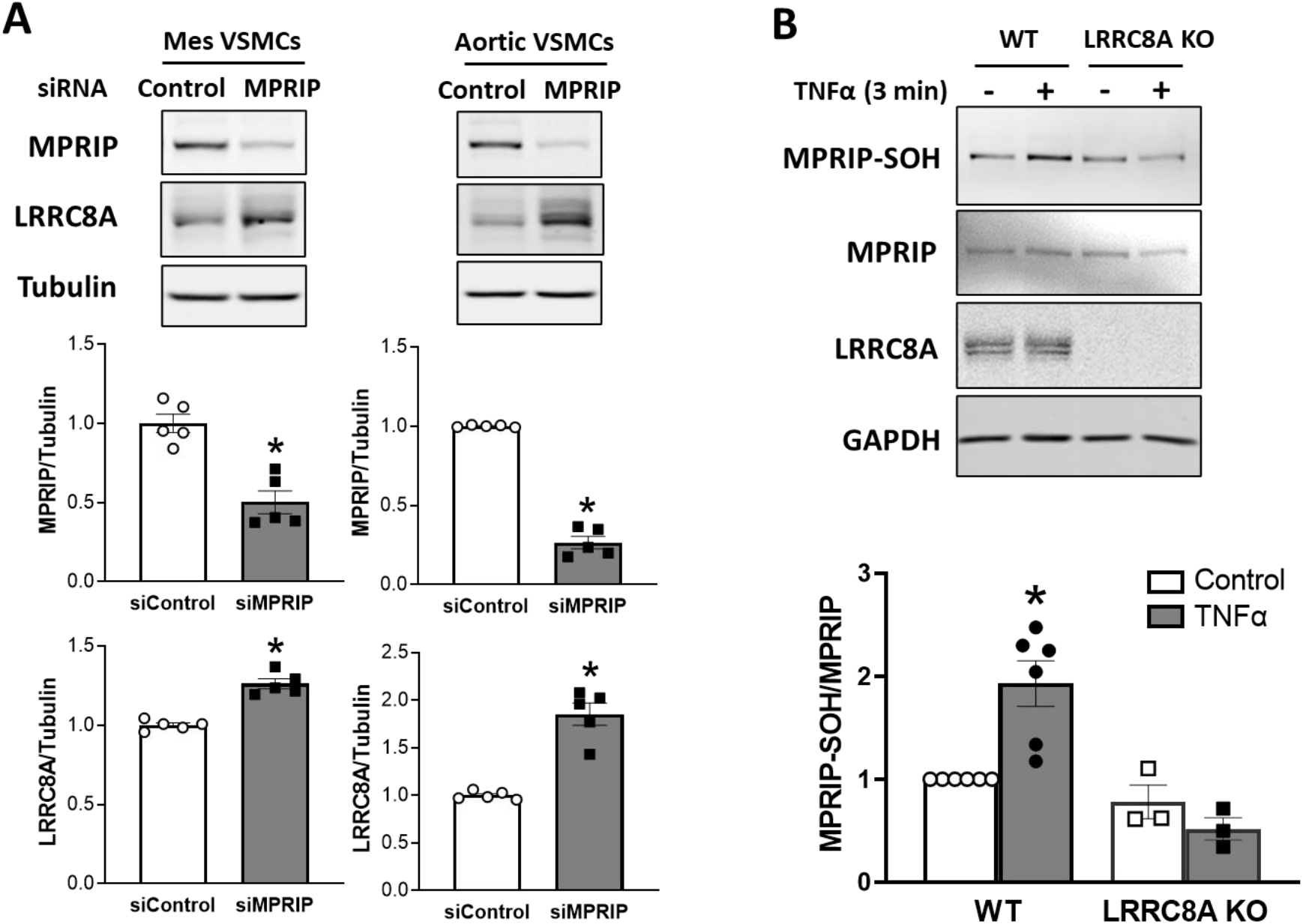
MPRIP knockdown increases LRRC8A expression and MPRIP is sulfenylated in response to TNFα. **A)** siRNA knockdown of MPRIP in cultured mesenteric (Mes) or aortic VSMCs resulted in a significant decrement in the abundance of MPRIP protein. In both cells this was associated with a significant increase in LRRC8A protein. **B)** A 3 min exposure to TNFα induced sulfenylation of MPRIP in WT but not KO VSMCs. The top panels provide representative blots, bar graphs summarize the results of 3 - 6 independent experiments.

The Oxymouse database provides quantitative mapping of the mouse cysteine redox proteome (https://oximouse.hms.harvard.edu). It defines 8 redox-susceptible cysteines in murine P, including sites within the binding regions for all 4 of its partner proteins (Fig 5D). We hypothesized that association with LRRC8A places of MPRIP in a nanodomain that undergoes significant redox shifts in response to TNFα. We assessed the redox status of MPRIP in live cells by quantification of cysteine sulfenylation with DCP-Bio1 under control conditions and after exposure to TNFα for 3 min. MPRIP was sulfenylated at rest and this increased significantly after TNFα exposure in WT, but not in KO VSMCs. Thus, TNFα causes MPRIP oxidation in an LRRC8A-dependent manner. Sulfenylation can alter protein structure and may affect the ability of MPRIP to associate with its binding partners. This may be an important link in the chain of redox-driven events that link TNFα receptor occupation to changes in VSMC contractility.

## Discussion

We previously demonstrated that LRRC8A VRACs are required for Nox1 activation by TNFα^16^. Here we identify LRRC8A channels as membrane-spanning links in a multi-protein complex that connects Nox1 to the actin cytoskeleton. While VSMC-specific LRRC8A knockout vessels displayed normal contractile responsiveness to depolarization (KCl) or alpha-adrenergic receptor activation (PE), they completely lacked myogenic tone. Vasodilation to ACh and SNP was augmented, and consistent with the requirement of LRRC8A for signaling, KO vessels were completely protected from TNFα-induced impairment of dilation. Identification of cytoplasmic proteins that associate with LRRC8A revealed association with the actin cytoskeleton. Binding of LRRC8A to MPRIP places it in proximity to RhoA, the MLCP complex and actin, critical regulators of contraction. RhoA activity was reduced by knockdown (siLRRC8A) or inhibition (CBX) of VRACs, as was ROCK-dependent phosphorylation of MYPT1 in KO mesenteric vessels. Collectively, these data suggest that decreased RhoA/ROCK activity contributes to enhanced relaxation and loss of myogenic tone. Importantly, MPRIP becomes oxidized following TNFα exposure, raising the possibility of another level of regulation; dynamic changes in MPRIP association with its partner proteins. While the details of how this multi-protein complex is assembled and regulated remain to be determined, we propose that association of Nox1/LRRC8A with MPRIP colocalizes ROS production with RhoA/ROCK and MYPT1, providing a mechanism by which Nox1-derived superoxide modulates calcium sensitivity.

Nox1 plays a critical role in inflammatory vasculopathy based on the ability of Nox1 knockout to rescue vascular dysfunction in multiple disease states (reviewed in^31^). VRACs support Nox1 activation, as evidenced by the fact that knockdown of LRRC8A or LRRC8C reduced Nox1-dependent extracellular O_2_^-•^ production and multiple downstream effects of TNFα^16,17^. LRRC8 channels share significant structural similarity with connexins and both channels are inhibited by CBX^32,33^. Connexins provide a pathway for conducted hyperpolarization from endothelial cells to VSMCs^34^ and blocking them impairs endothelium dependent VSMC hyperpolarization in response to ACh^32^. The concentration of CBX used (100 μM) was shown by others to produce ~75% block of VRAC^30^. In endothelium-intact rat hepatic and mesenteric arteries stimulated with 1 μM PE, this concentration of CBX caused a small depolarization of VSMCs but had no effect on ACh-induced hyperpolarization in the hepatic artery and caused only very mild inhibition in the mesenteric^32^. This suggests that the majority of voltage-dependent relaxation in these vessels is mediated by activation of K^+^ channels by endothelial-derived factors^35^, rather than by conducted hyperpolarization from the endothelium. This contribution is demonstrated by the partial inhibition of ACh-induced relaxation by apamin + charybdotoxin. However, control and CBX-treated vessels were similarly impacted based on similar degrees of rightward shift in the dose-response relationship (Fig. 1). This suggests that the ability of LRRC8A KO and CBX to enhance relaxation is related to a mechanism that is independent of changes in membrane voltage and associated alterations in calcium influx.

As hypothesized, KO vessels were protected from TNFα-induced injury, but the virtually complete protection that was observed was unanticipated as TNFα in known to cause both VSMC and endothelial dysfunction^36^. These data suggest that endothelial injury induced by TNFα in this *in vitro* model was minimal and impaired relaxation was almost completely a function of VSMC dysfunction. This interpretation is supported by the fact that responses to ACh and SNP were similarly affected. The ability of short-term exposure to CBX to restore both ACh and SNP responsiveness following a 48hr exposure to TNFα was remarkable (Fig. 2). The similarity between the impact of LRRC8A KO and VRAC inhibition suggests that impairment of relaxation induced by TNFα is dependent upon VRAC function and caused by a sustained process that does not occur in the KO vessels.

Increased relaxation to ACh and SNP could be related to enhanced NO^•^ responsiveness, and/or to an increase in NO^•^ bioavailability. The latter might be the result of a decrease in O_2_^-•^ production by Nox1 when LRRC8A is absent or VRACs are blocked^16^. However, this mechanism was not supported by the persistence of enhanced vasodilation when proximal steps in NO^•^ signaling were bypassed and PKG was directly activated by BAY60-2770. Enhanced dilation was also not uniquely a property of PKG-dependent signaling as relaxation in response to PKA stimulation with forskolin, or ROCK inhibition by Y-27632 were also potentiated (Suppl Fig. 2). These data point to an impact of VRAC on a process that is downstream of PKA, PKG and RhoA/ROCK. These kinases all control MLCP activity via phosphorylation of MYPT1.

NADPH oxidases play critical roles in vascular mechanosensing^37^. Spontaneous tone in rat cremasteric and cerebral was reversibly blocked by inhibition of Nox1^19^ Pressure-induced renal afferent arteriolar tone is associated with a burst of oxidant production that is NADPH oxidase-dependent and is inhibited by disruption of O_2_^-•^ (apocynin, diphenylene iodonium, PEG-Superoxide Dismutase, Tempol) but not H_2_O_2_ (PEG-catalase) bioavailability^38^. Pressurization of mouse mesenteric arteries induced MYPT1 phosphorylation at T853, and myogenic tone was completely blocked by Y-27632^20^. Loss of myogenic tone in LRRC8A KO mesenteric arteries (Fig. 3) is thus consistent with disruption of a Nox1/LRRC8A/O_2_^-•^/RhoA/ROCK signaling axis. In endothelial cells, loss of LRRC8A impaired Akt-dependent eNOS activation by stretch and disrupted the ability of cells to orient to shear stress^39^. Importantly, O_2_^-•^ produced by endothelial Nox2 activates Akt in response to shear stress^40^ and Nox2 also physically associates with LRRC8A^41^. Future work will be required to sort out the relative contribution of VRACs to endothelial and VSMC dysfunction in pathologic states.

The proteins pulled down by LRRC8A strongly support an association with the actin cytoskeleton. MPRIP was chosen for further investigation as a direct binding partner due to its established connection with proteins that are critical to the regulation of contractility, including actin. The two proteins share a similar pattern of plasma membrane distribution with a particularly strong association in regions of membrane ruffling (Fig. 5C). Importantly, this is where the highest levels of H_2_O_2_ were detected by a dynamic, actin-associated sensor^42^. Colocalization of any protein with Nox1/LRRC8A raises the possibility that this facilitates redox regulation. For instance, two unconventional myosins (Myo1c and Myo1d) associated with LRRC8A. Myo1d binds to and localizes epidermal growth factor receptors (EGFRs) to the plasma membrane^43^. EGFR transactivation occurs in response to AngII in an NADPH oxidase and ROS-dependent manner and inhibition of Nox oxidases normalized increased EGFR phosphorylation in spontaneously hypertensive rats^44^.

RhoA activation regulates cytosolic calcium concentration by multiple mechanisms including regulation of calcium and potassium channels^11^. Reduced RhoA activity induced by siLRRC8A or CBX (Fig. 6A) is consistent with enhanced relaxation. The relatively modest degree of impairment may be due to the fact that RhoA fulfills other roles beyond regulation of ROCK or could point to the involvement of other mechanisms. RhoA activity is enhanced by oxidation of specific cysteine residues in its active site which enhances the dissociation rate of GDP^45^. In contrast, over-oxidation of these residues can cause formation of an intramolecular disulfide that inhibits RhoA activity^46^. In isolated vascular rings, extracellular generation of O_2_^-•^ using a mixture of xanthine and xanthine oxidase induced vasoconstriction that was blocked by the SOD mimetic tempol, but not by catalase. These responses were associated with an increase in MYPT1 phosphorylation by ROCK^47^. This suggests that RhoA can be activated by extracellular O_2_^-•^ which supports the concept of O_2_^-•^ influx but does not preclude a role for cytoplasmic H_2_O_2_ that is formed after O_2_^-•^ enters the cell.

MYPT1, the regulatory subunit of MLCP, is the master regulator of calcium sensitivity. It controls activity of the PP1C subunit of the MLCP complex. This phosphatase counteracts MLC kinase (MLCK) mediated, calcium-calmodulin dependent phosphorylation of the 20kD subunit of MLC. MYPT1 can thereby modulate the rate of actin-myosin crossbridge cycling at a constant cytoplasmic calcium concentration. MYPT1 is regulated by multiple kinases. ROCK inhibits MYPT1 by phosphorylation at both T696 and T853. Directly adjacent to these sites are serine residues (S695, S852) that are targets for activation by PKA and PKG. Phosphorylation at the ROCK sites is inhibited by the presence of a phosphate at the PKG and PKA sites and *vice versa*, creating interdependence of MLCP activation and inhibition. Reduced MYPT1 phosphorylation at T853 in LRRC8A KO mesenteries is consistent with a phenotype of enhanced vasodilation, but this may not provide a complete explanation for the functional changes observed when LRRC8A is absent.

MPRIP targets RhoA to stress fibers and facilitates dephosphorylation of myosin by MLCP. Silencing of MPRIP prevented RhoA/ROCK-dependent phosphorylation of MYPT1 but did not interfere with activation of RhoA and ROCK in A7r5 VSMCs^26^. Association of MPRIP with MYPT1 enhances phosphatase activity^48^ while binding to RhoA is inhibitory^49^. These findings point to an important role for MPRIP complex assembly. Thus, enhanced dilation in KO vessels may be related to a combination of reduced RhoA activity and a change in the association of proteins with MPRIP. MPRIP oxidation in response to TNFα (Fig. 7B) adds functional evidence of association with Nox1/LRRC8A and raises that possibility that MPRIP binding to its scaffolded partners is oxidation-dependent. A significant contribution of complex assembly may explain why relaxation to SNP was augmented in LRRC8A null mice but was not altered in vessels from Nox1 null mice^50^. Future work will determine how redox signals impact MPRIP complex assembly and vasomotor function.

When LRRC8A is missing or VRACs are blocked, extracellular O_2_^-•^ production by Nox1 in response to TNFα is reduced^16^. We have proposed that anion flux through LRRC8A provides charge compensation for electron flow through Nox1 and speculated that VRACs may also serve as a O_2_^-•^ conductance allowing extracellular O_2_^-•^ to come back across the plasma membrane^6^. While it is not possible to accurately quantify the forces acting on O_2_^-•^ immediately following its creation, the initial essentially infinite concentration gradient clearly favors O_2_^-•^ influx, but the concentration of O_2_^-•^ is very low. However, current flow through NADPH oxidases produces significant membrane depolarization^51^. This local event decrements at increasing distance from the oxidase but its potential to impact O_2_^-•^ influx is maximized by physical association of LRRC8A with Nox1. The concept of a O_2_^-•^ flux through an anion channel is supported by detection of O_2_^-•^ efflux from endosomes that is sensitive to anion channel blockers^52^. Cytoplasmic dihydroethidium oxidation by extracellular O_2_^-•^ was measured and inhibited by anion channel blockade or knockdown of Chloride Channel three (ClC-3) expression^53^. We recently demonstrated that ClC-3 influences membrane trafficking of LRRC8A, raising the possibility that this effect of ClC-3 was indirect^27^.

## Summary and significance

We propose that association of LRRC8A with MPRIP provides a critical, membrane-spanning link in TNFα signaling. A proposed model of Nox1-dependent redox signaling (Fig. 8) incorporates MPRIP interaction with all four of its partner proteins. Extracellular O_2_^-•^ is created in proximity to LRRC8A VRACs. Local depolarization could both activate LRRC8A and drive O_2_^-•^ influx. Of note, VRACs are also regulated by oxidation^17,54^ which may add an extra layer of regulation to the system. In the cytoplasm, O_2_^-•^ would be deposited into a H^+^ rich cytoplasmic nanodomain created by Nox1-dependent NADPH metabolism^55^. These protons could support local creation of H_2_O_2_ as O_2_^-•^ dismutation is favored by acidic conditions (pKa of H_2_O_2_ = 11.75). Association with LRRC8A places MPRIP and RhoA in this oxidized nanodomain. ROCK activation may be disrupted when LRRC8A is absent or blocked. Reduced ROCK activity favors MYPT1 phosphorylation by PKG or PKA and potentiates relaxation. The scaffolding function of MPRIP may be redox-dependent, allowing physical interaction to impact MYPT1 and therefore MLCP activity.

**Figure 8.**
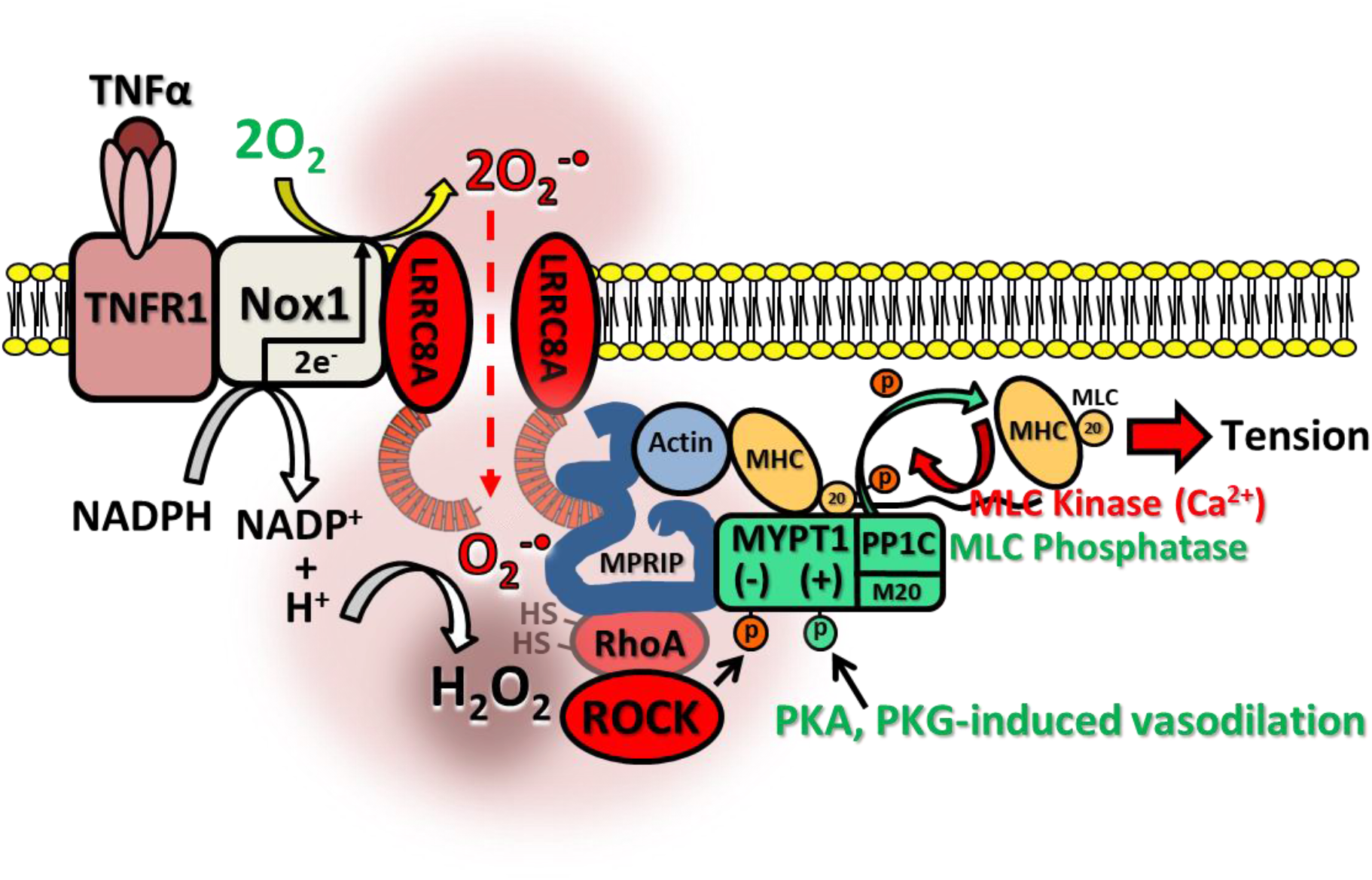
TNFR1/Nox1-dependent regulation of VSMC contractility. TNFR1 associates with Nox1 and LRRC8A at the plasma membrane. TNFα activates Nox1, resulting in NADPH conversion to NADP^+^ + H^+^ and 2 electrons are passed across the membrane and donated to O_2_ to produce extracellular O_2_^-•^. Local membrane depolarization by Nox1 facilitates O_2_^-•^ influx through LRRC8A into a proton rich environment which may facilitate localized H_2_O_2_ production, contributing to creation of an oxidized nanodomain. These oxidants promote redox cycling of RhoA, causing ROCK activation and inhibitory phosphorylation of MYPT1. This reduces myosin light chain phosphatase (MLCP = MYPT1 + PP1C + M20) activity, increasing phosphorylation of the 20kD subunit of MLC and enhancing the calcium sensitivity of MLC which promotes contraction. MPRIP is also oxidized which may impact complex assembly. Loss or inhibition of VRACs inhibits Nox1 activity and prevents O_2_^-•^ influx, reducing cytoplasmic oxidation, RhoA activation and ROCK phosphorylation of MYPT1. This enhances the ability of PKA and PKG to phosphorylate MYPT1, activate MLCP, and dephosphorylate MLC, thus reducing calcium sensitivity and potentiating VSMC relaxation.

Hypertension is a chronic inflammatory disease that is associated with enhanced vascular contractility^56^. TNFα^36,57^ and AngII^3,58^ signaling are increased and promote cardiovascular damage. Both pathways depend upon Nox1 activation^5–7,59^ and are associated with “oxidative stress”. This work provides a mechanism for Nox1-dependent redox signaling that utilizes LRRC8A anion channels to link extracellular O_2_^-•^ to RhoA activation and MLCP inhibition. This may allow inflammatory signaling via ROS to alter vasomotor function.

## Sources of Funding

This work was supported by NIH R01 HL160975 and R01 DK132948

## Notes

### Competing Interest Statement

The authors have declared no competing interest.

